# DNA transposons of *maT* family in the Cnidaria

**DOI:** 10.1101/2022.10.13.512200

**Authors:** Mikhail V. Puzakov, Lyudmila V. Puzakova, Sergey V. Cheresiz, Shasha Shi

## Abstract

Transposable elements exert a significant influence on the structure and size of eukaryotic genomes. Representatives of *Tc1/mariner* superfamily of DNA transposons form a prevalent and highly variable group, which includes the relatively well studied *TLE*/DD34-38E, *MLE*/DD34D, *maT*/DD37D, *Visitor*/DD41D, *Guest*/DD39D, *mosquito*/DD37E and *L18*/DD37E families. A detailed study of distribution and diversity of *Tc1/mariner* transposons will help us to better investigate the co-evolution of TEs and eukaryotic genomes. We performed a profound analysis of *maT*/DD37D family in the cnidarians. maT transposons were shown to exist in a limited number of cnidarian species belonging to Cubozoa, Hydrozoa and Scyphozoa classes. *maT* transposons of the cnidarians are thought to be the descendants of several individual invasion events, which have occurred at different times in the past. The *mosquito*/DD37E transposons of the cnidarians have also been described. These TEs were shown to be present in Hydridae family (class Hydrozoa) only. An analysis of TE distribution, diversity, evolutionary history and phylogeny established that theTEs undergo their unique evolution not only in different species, but also within a particular species. These results improve our knowledge of *Tc1/mariner* diversity and evolution, as well as their influence of eukaryotic genomes.

## Introduction

Transposable elements (TE) represent the DNA fragments without fixed localization, which are able to transpose within the host genome. TEs have been identified, virtually, in every genome, from bacteria to humans. These DNA fragments can considerably increase their copy numbers, while causing changes in genome size, structure and function (Casacuberta and González 2013; Sotero-Caio et al. 2017; Bourque et al. 2018; Arkhipova and Yushenova 2019). TEs can also become “domesticated” and serve as a source for new host genes, thus, taking part in the adaptation and evolution processes (Casacuberta and González 2013; Sotero-Caio et al. 2017; Bourque et al. 2018).

Mobile elements of eukaryotes are represented by two different classes of retrotransposons and DNA transposons. This classification is based on the mechanisms of transposition of mobile elements. The motility of retrotransposons occur via the activity of reverse transcriptase, and the RNA intermediate, the newly synthesized DNA copy of which is integrated into the genome. This process is referred to as the copy-and-paste mechanism. The motility of DNA transposons does not require an intermediate, since they use transposase to cut out of the previous locus and insert into a new one, the latter mechanism being referred to as “copy-and-paste” (Wicker et al. 2007; Kapitonov and Jurka 2008; Kojima 2020).

The *Tc1/mariner* superfamily is one of the most common and prevalent ones among the DNA transposons (Feschotte and Pritham 2007). It includes a number of families, such as the *Tc1*-like (*TLE*/DD34-38E) and mariner-like (*MLE*/DD34D) elements, as well, as *maT*/DD37D, *Visitor*/DD41D, *Guest*/DD39D, *mosquito*/DD37E and *L18*/DD37E, (Shao and Tu 2001; Tellier et al. 2015; Zhang et al. 2016; Dupeyron et al. 2020; Shen et al. 2020; Wang et al. 2021; Puzakov and Puzakova 2022). *maT* family has been first described in 2002. The authors of this publication demonstrated that *maT* represents an intermediate group between *Mariner* and *Tc1* families (Claudianos et al. 2002). The size of an intact representative of this family is about 1.3 thousand bp. They contain an ORF coding for the transposase enzyme, which consists of around 346 amino acid residues. The transposase gene is flanked by the terminal inverted repeats (TIRs), the size of which ranges between 13 and 48 bp. (Wang et al. 2021). The other structural features of *maT* transposase, which are shared by the representatives of *Tc1/mariner* superfamily, have also been described. PAIRED domain involved in DNA binding is localized to N-terminal part of transposase sequence (Nagy et al. 2004; Rousseau 2004). The DNA-binding domain can be readily recognized due to the secondary structure of transposase with 3 alpha-helices representing each of its subdomains, PAI and RED, which are divided by a short spacer. In addition to PAIRED domain, GRPR motif located between PAI and RED subdomains is another common feature of *maT* transposase. Most of *maT* transposases possess a nuclear localization signal (NLS). The catalytic domain of *maT* transposons has a specific pattern, DD37D, with a spacer between second and third residues of aspartic acid being 37 amino acid residues in length. This domain localized to the C-terminal part of transposase is involved in the cut-and-paste mechanism (Ivics et al. 1997; Plasterk et al. 1999; Ivics and Izsvák 2015; Tellier et al. 2015; Wang et al. 2021).

*maT* family is rather understudied despite the fact that it has been described as long ago, as in 2002. In their 2021 publication, Wang and co-authors reported that *maT* transposons were found mainly in invertebrates. According to the results of phylogenetic analysis, they classified *maT* transposases into 5 main clusters (A – E) (Wang et al. 2021).

In the present work, we study *maT* DNA transposons of the cnidarians (*Cnidaria*). The cnidarians belong to the type of multicellular organisms, which possess the stinging cells. These ancient orgamisms, which inhabit, exclusively, the water reservoirs (Technau and Steele 2011)., have emerged about 741 million years ago (http://www.timetree.org). A detailed study of the structure and diversity of *maT* elements of the cnidarians, will extend our knowledge of this transposon family, help us better understand the evolutionary dynamics and evolutionary history of these elements.

## Materials and methods

### Search for DD37D transposons

We searched for *maT* (DD37D) transposons using tBLASTn software with standard settings (https://blast.ncbi.nlm.nih.gov/Blast.cgi). Amino acid sequences of transposases of *maT-Hyvu, maT-Hyvi*, and *maT-Hyol* elements identified in *Hydra vulgaris, Hydra viridissima*, and *Hydra oligactis*, respectively (Wang et al. 2021), were used as a query. The complete genomic sequences of the representatives of the Cnidaria used in this research were extracted from NCBI database (https://www.ncbi.nlm.nih.gov). The search for the complete copies of mobile elements was performed using their homology to the above *maT-Hyvu, maT-Hyvi*, and *maT-Hyol* elements. The identified homologous sequences were analyzed with their flanking regions from the respective scaffolds spanning ~3000 bp. The sequences of the identified complete *maT* elements were further employed for the subsequent identification of their copies and the calculation of their copy numbers. The element copies were considered the complete ones when they possessed the transposase ORF longer than 300 amino acid residues and both terminal inverted repeats (TIRs). The elements, which were shorter, than 10% of the complete element’s size, were not taken into our analysis. The elements possessing from 10 to 100% of the complete element’s sequence were calculated as the total number of elements. Only those copies possessing 95-100% of the complete genomic sequence of an element were considered (and counted) as the full-genomic copies.

### Analysis of the transposon structures

We defined the boundaries of presumptive ORFs using ORF Finder software (https://www.ncbi.nlm.nih.gov/orffinder/), then, the ORF ends were refined visually. BLASTn software was used for TIR identification. An analysis of secondary structures with the use of PSIPRED v4.0 (Buchan and Jones 2019) enabled us to identify PAIRED motif. The catalytic (DDD) domain and GPRP-like motif were identified visually. Nuclear localization sequences (NLS) were identified using PSORT software (https://www.genscript.com).

### Phylogenetic analysis

Phylogenetic analysis was performed with the use of MEGA X software (Kumar et al. 2018). Our analysis included the amino acid sequences of previously described *Tc1/mariner* superfamily transposases and those of the cnidarians, which have been identified in the present work. The *IS630* transposases were used as an outgroup (see Supplementary materials). Multiple alignments of amino acid sequences was performed with the use of MUSCLE (Edgar 2004) with standard settings. The maximum likelihood method was employed for the calculation and construction of phylogenetic trees. The reliabilities of the topologies were evaluated using bootstrap method (1000 replications).

## Results and discussion

### maT transposons of the cnidarians

In order to search for DD37D transposons of the cnidarians, we used the amino acid sequences of transposases of several elements (*maT-Hyvu, maT-Hyvi*, and *maT-Hyol*) found in *Hydra vulgaris, Hydra viridissima*, and *Hydra oligactis*, respectively (Wang et al. 2021), as queries. Among the results obtained with the use of tBLASTn, we selected those sequences showing the highest degree of identity. An analysis of 77 complete genomic sequences of the cnidarians (Fig. 1) yielded 19 *maT* elements identified in 9 species. These organisms belong to 3 classes of the cnidarians (Cubozoa, Hydrozoa и Scyphozoa), while 3 other classes (Anthozoa, Myxozoa, and Staurozoa) were lacking *maT* elements (Fig. 1). Only ten out of nineteen elements identified possessed the complete copies, while the rest of the elements were either truncated copies or those lacking the TIRs (Supplemental Material 1). The complete elements were 1292-1602 bp in size, with their transposases possessing 320-350 amino acid residues, both features being typical for this family. Twelve out of nineteen elements possessed the TIRs, 18-139 bp in length (Supplemental Material 1). The latter feature is different from the typical TIR length of the previously reported *maT* elements (13-48 bp) (Wang et al. 2021). However, the atypically long TIRs (69-139 bp) were found in 4 out of 12 elements, only, while the rest eight had the TIR size ranging from 18 to 39 bp, which is a typical feature of *maT* elements. In addition to TIRs, *maT-1_CFus* element had the subterminal inverted repeats (SIRs) 38/39 bp long, while *maT-1_HOli* element had only one terminal repeat (Supplemental Material 1). SIRs are found in some representatives of *Tc1/mariner* superfamily (Zhang et al. 2016; Puzakov et al. 2021). The sizes of truncated *maT* copies ranged from 216 to 1575 bp, with the exception of *maT-4_HVir/GCA_014706445*, which had the size of 3159 bp (Supplemental Material 1) due to a long noncoding insertion breaking the ORF of this element.

**Fig. 1.**
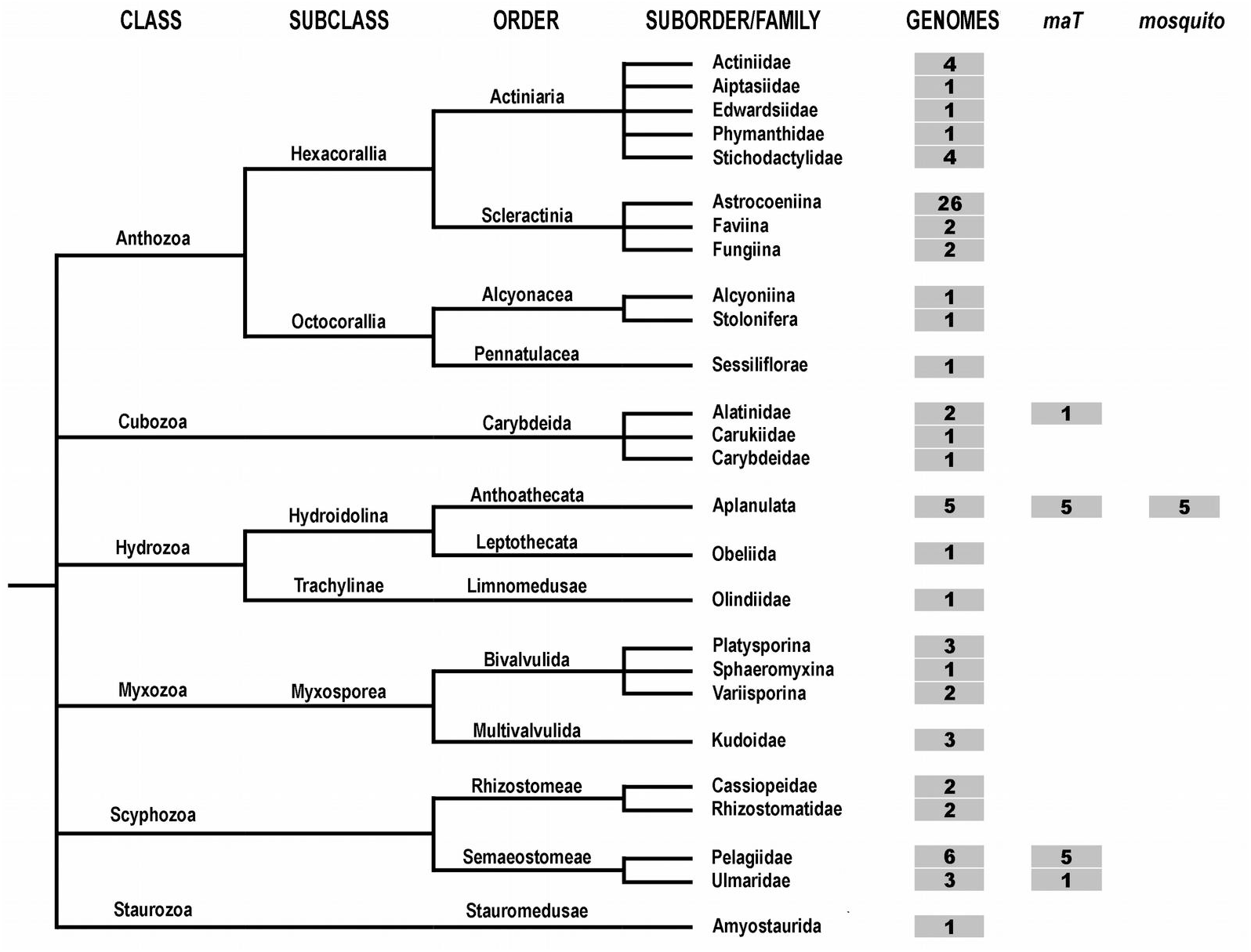
Prevalence of *maT* and *mosquito* elements in the cnidarians. Only the groups of organisms possessing the sequenced genomes are represented in the taxonomic tree. The columns (in gray boxes) indicate the number of whole genome sequences available for analysis and the number of genomes in which the corresponding groups of DNA transposons were found.

An analysis of *maT* copy numbers showed that 4 of the described 19 elements had a low copy number varying from 2 to 13. Nine elements had moderately high copy numbers (27-83), while the rest six were represented by multiple copy numbers (162-376) (Supplemental Material 1). Also, four elements possessed considerably high numbers of complete copies (24-46), one element had 11, while five had 3-5 complete copies (Supplemental Material 1). Nine *maT* elements were lacking the complete copies.

### An analysis of transposase structures of *maT* transposons

In order to identify the potentially functional transposons among the complete *maT* elements, we selected the group of transposons with an intact ORF, since any breaks (frameshifts) or stop codons in the ORF will disable the encoded transposases. Out of 10 transposons possessing the complete copies, only 7 met the requirements for the above selected group. The transposase amino acid sequences of two elements, *maT-1_CQui/GCA_012295145* and *maT-1_CQui/GCA_014526335*, turned out to be completely identical in both assemblies of *Chrysaora quinquecirrha* genome, thus, only one copy was taken into the analysis. The analysis of PAIRED domain playing an important role in DNA binding (Nagy et al., 2004; Rousseau 2004; Zong et al., 2020) showed that first three alpha helices of PAI subdomain were present in 5 *maT* elements, only (Fig. 2). In *maT-1_CQui/GCA_012295145* element, the amino acid sequence of transposase lacks the full PAI subdomain. Conversely, the three alpha helices of RED subdomain are present in all elements. However, the third alpha helix of *maT-2_HVir_GCA_014706445, maT-1_HVir_GCA_004118115*, and *maT-3_HVir_GCA_004118115* elements is shortened, while it is fragmented in *maT-1_CQui/GCA_012295145* element (Fig. 2).

**Fig. 2.**
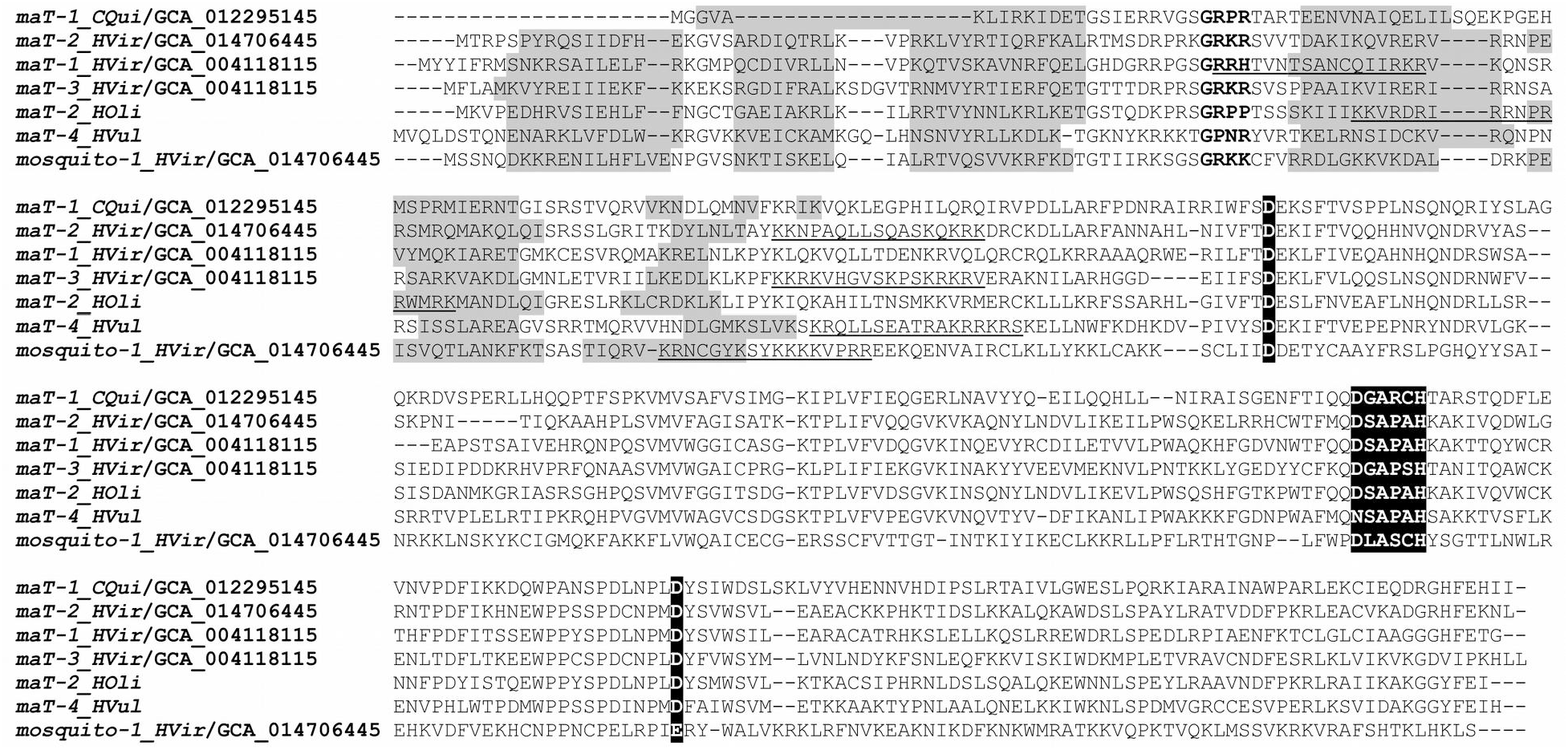
Multiple alignment of the amino acid sequences of *maT* and *mosquito* transposases. Six α-helices of PAIRED domain are highlighted gray; a hypothetical nuclear localization signal (NLS) is underlined; the GRPR-like motif is in bold; marker loci of the DDE/D domain are highlighted black.

GPRP-like motif localizing between PAI and RED subdomains was found in any transposase of the analyzed *maT* elements (Fig. 2). NLS was identified in the transposases of five elements, although its localization in *maT-1_HVir_GCA_004118115* and *maT-2_HOli* transposases was not rather typical (Fig. 2). NLS localization between DNA-binding and catalytic domains is typical for active *Tc1/mariner*-transposons (Plasterk et al. 1999). Meanwhile, NLS was not found in *maT-1_CQui/GCA_012295145* element (Fig. 2). The catalytic DD37D domain was identified in all complete *maT* elements studied (Fig. 2). However, the second aspartate residue (D) of DD37D domain is substituted for asparagines (N), which, likely, disabled its function, since a fully conserved catalytic domain is indispensable for the cut-and-paste function of the transposon (Ivics and Izsvák 2015). Summarizing all above data, four *maT* transposons (*maT-2_HVir/GCA_014706445, maT-1_HVir/GCA_004118115, maT-3_HVir/GCA_004118115* and *maT-2_HOli*) can be potentially functional, since they possess both TIRs and an intact transposase with all domains required for the transposition.

### Phylogenetic analysis of *maT* elements

The phylogenetic analysis of all *maT* transposones identified in the cnidarians shows that they form a single group with the other *maT* transposones included in our analysis. However, this clade has a low bootstrap value (below 50%) (Fig. 3), which may indicate that this family is not monophyletic, but it, rather, joins several phylogenetically distinct groups. The latter is contrary to the previous reports showing that this family forms a single branch with high confidence (Dupeyron et al. 2020; Wang et al. 2021).

**Fig. 3.**
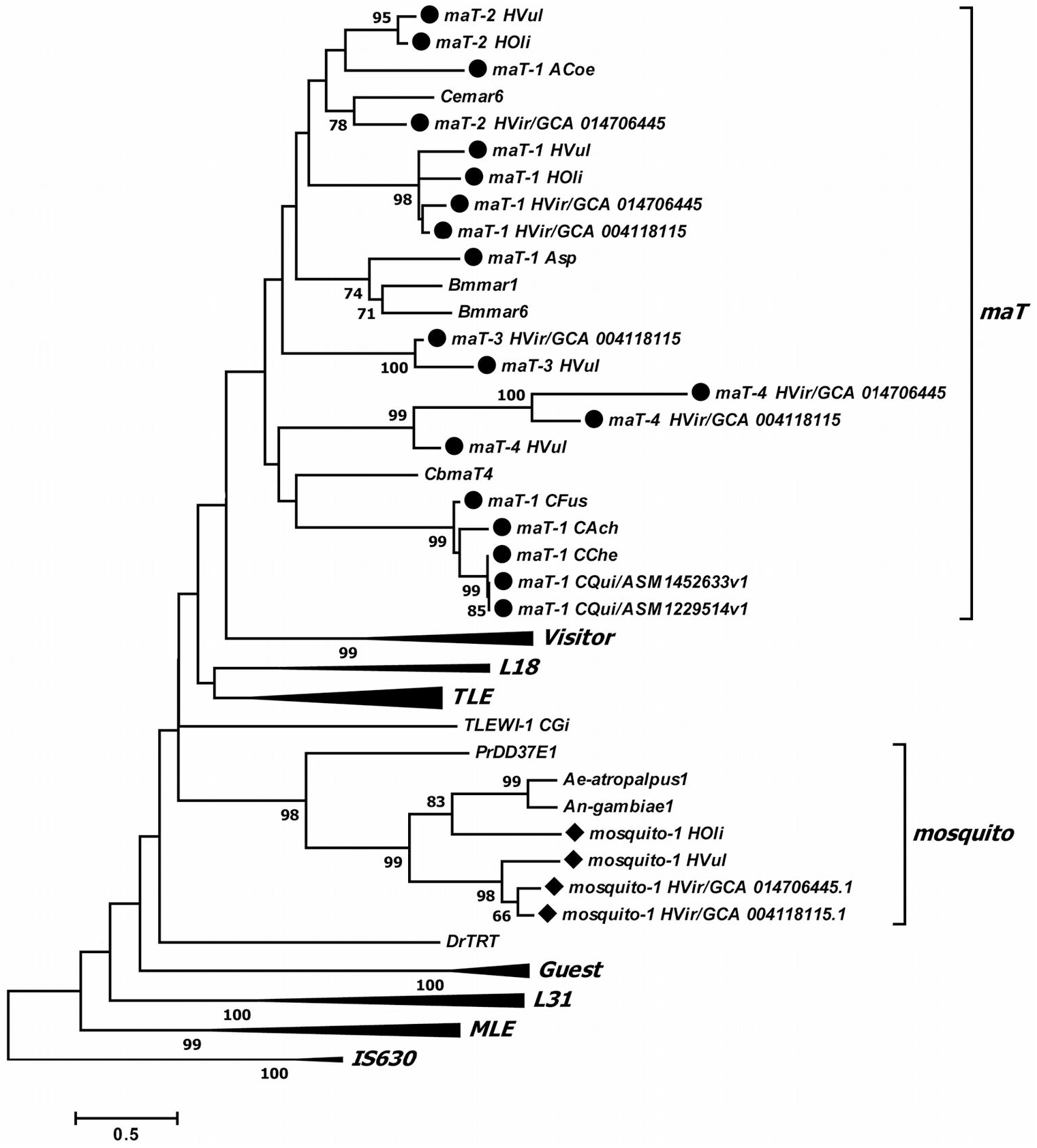
Evolutionary relationships between the *maT, mosquito* and other *Tc1/mariner* transposons. Black circles (*maT*) and rhombuses (*mosquito*) indicate DNA transposons found in this work. Phylogenetic analysis was performed using the MEGA X software with the maximum likelihood method. The WAG+G+F model was used. The reliability of the topology was assessed using the bootstrap test (1000 replications). Bootstrap value below 50% are not shown.

To study the phylogenetic relationships within *maT* family more accurately, we analyzed all the transposases of *maT* elements discovered in our work with those of the other *maT* elements published by Wang and co-authors (Wang et al. 2021). In this publication, all *maT* elements were classified into five clusters designated by letters A to E. Several complete copies of *maT* transposons were selected from each group for the phylogenetic which showed some variation from the original classification (Fig. 4). According to our analysis, elements from clusters A and B split into the groups, which were located distantly on the phylogenetic tree. Thus, A and B clusters became split into A1 and A2, and B1 and B2, respectively. The names to these new clusters were given according to the original names provided by Wang and collaborators (Wang et al. 2021). Fourteen out of the described 19 *maT* elements of the cnidarians were classified into 4 clusters (from A to D), including the subgroups (Fig. 4). Five elements belonging to *Pelagiidae* family formed a new cluster with high bootstrap value, which we designated as F. None of *maT* elements identified in our work was classified into E cluster. Also of note, not every cluster demonstrates the high bootstrap value for all its members.

**Fig. 4.**
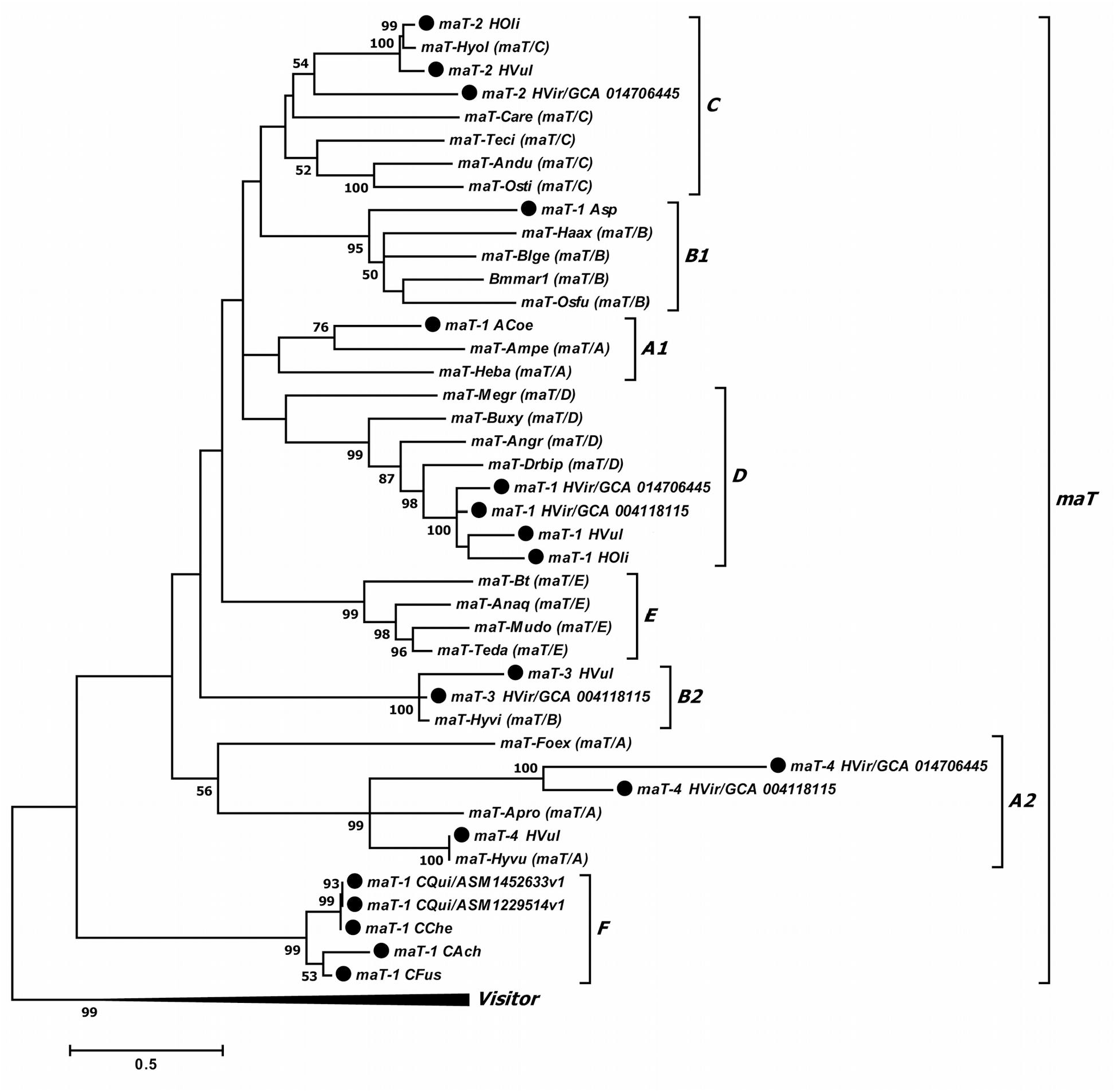
Phylogenetic tree of *maT* elements. Black circles indicate DNA transposons found in this work. Phylogenetic analysis was performed using the MEGA X software with the maximum likelihood method. The LG + G model was used. The reliability of the topology was assessed using the bootstrap test (1000 replications). Bootstrap value below 50% are not shown.

### Evolutionary dynamics of mAT transposons

The time of insertion of a transposon into host genome can be generally estimated using the formula T = k/2r (Kimura 1980; Schemberger et al. 2019; Zong et al. 2020), where T is the time of insertion, k – is Kimura two-parameter distance, and r is the neutral mutation rate. The rate of synonimous substitutions in the representatives of Anthozoa is known to be 5 x 10^−10^/site/year, which is ~50-100-fold lower, than that of the other animals (Hellberg, 2006). Conversely, the rate of nucleotide substitutions in the hydrozoans and the scyphozoans is not that low according to the current scientific data (Dawson and Jacobs 2001; Govindarajan et al. 2005). Meanwhile, in few publications of some scholars, the time of insertion is estimated with the use of average substitution rate (10^−8^/site/year) (Schemberger et al. 2019). Since we could not find the accurate values of neutral mutation rates in the classes of Cubozoa, Hydrozoa and Scyphozoa, we did not recalculate the values of the statistical parameter k into particular temporal values when plotting.

Our estimates of the evolutionary dynamics of *maT* transposons of Scyphozoa (Fig. 5) yielded the values of the parameter k, which corresponded to the time of insertion of these elements into the host genomes and ranged within 34-42%. If we substitute the neutral mutation rates of 10^−8^/site/year into the formula for the calculation of insertion times, we will arrive to the time of invasion of maT transposons of the scyphozoans into the genome of their ancestral organism ranging from 21 to 17 MYA (millions years ago). Despite the copy numbers of *maT-1_CQui* elements in the genomes of different individuals of *Chrysaora quinquecirrha* being different, the evolutionary dynamics and the genetic coverage values of these elements were rather similar. The peak of activity of this element occurred far ago (around 8-10 MYA). The activity of *maT-1_CChe* element found in the genomes of *Chrysaora chesapeakei* occurred nearly in the same period, while no apparent period of TE activity was observed in *Chrysaora fuscescens*.

**Fig. 5.**
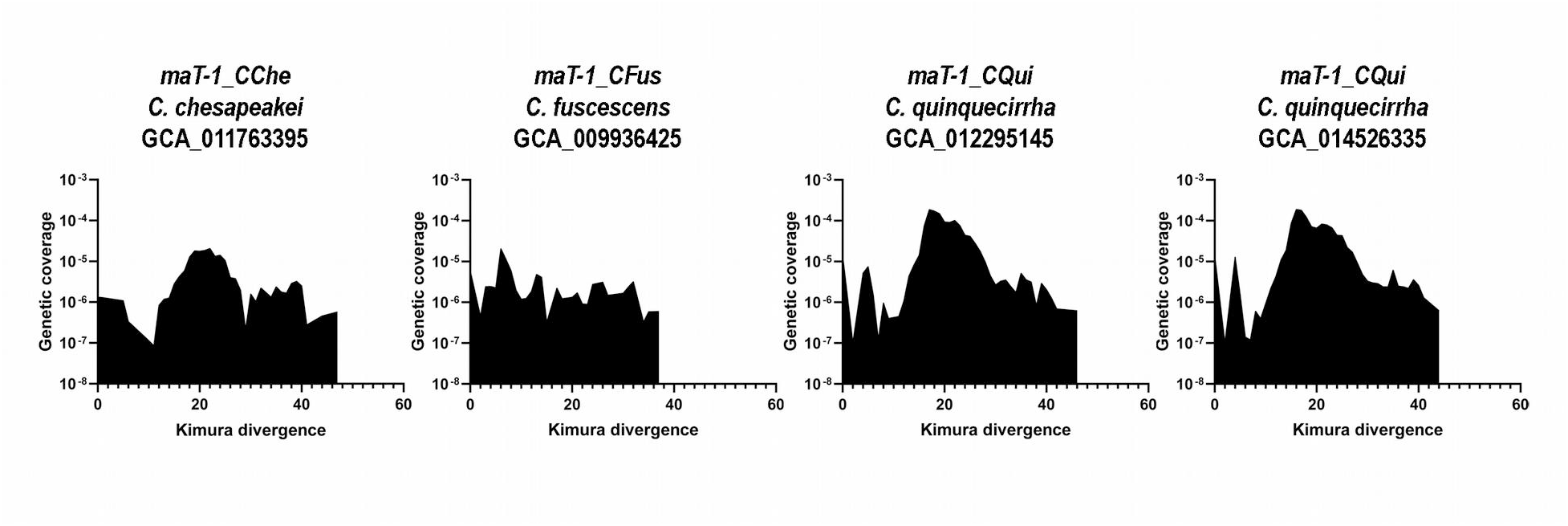
Evolutionary dynamics of cnidarian *maT* transposons. The X-axis indicates the Kimura divergence estimate (%), the Y-axis represents the log10 of copies coverage (%) in the genome.

We further studied the evolutionary dynamics of *maT* transposons of the hydrozoans. Since all four *maT* types (Supplemental Material 1) could not be identified in every representative of the hydrozoans, the genomes of those lacking some of *maT* types were analyzed using the sequence of a homologous TE present in another species of the hydras as a query. This was done in order to refine the *maT* search data obtained by tBLASTn. In our analysis, virtually, all elements of the hydrozoans also demonstrated high values of Kimura two-parameter distance (> 40%) and the estimated times of insertion > 20 MYA (provided the neutral mutation rate equals to 10^−8^/site/year). In addition, the RepeatMasker alignment algorithms revealed the homologies of those *maT* element types, which could not be discovered by tBLASTn (Fig. 6).

**Fig. 6.**
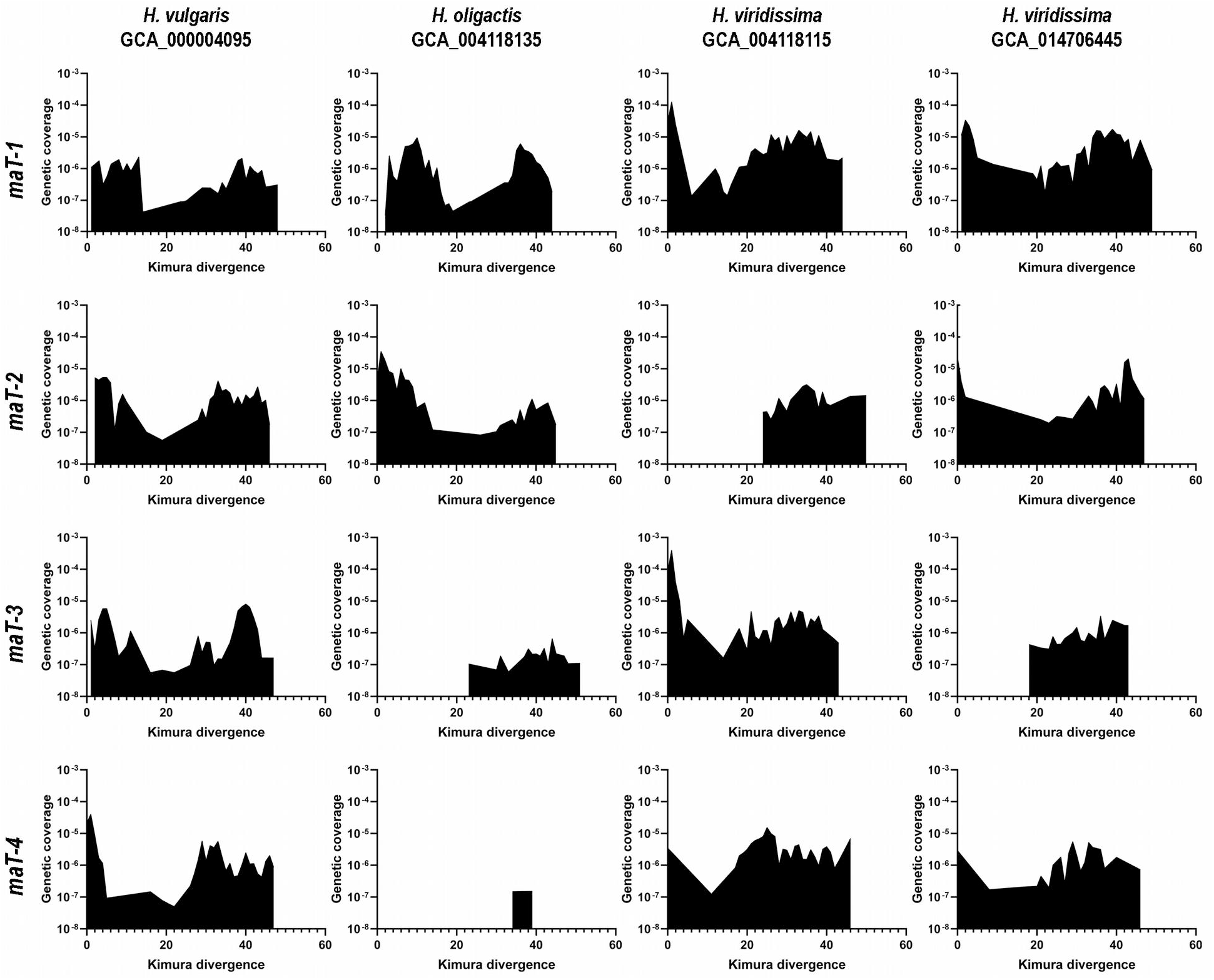
Evolutionary dynamics of hydras *maT* transposons. The X-axis indicates the Kimura divergence estimate (%), the Y-axis represents the log10 of copies coverage (%) in the genome.

The evolutionary dynamics plots constructed for *maT-1* elements, which are present in all genomic assemblies of the hydrozoans, demonstrate two activity peaks in every case. However, an analysis of both *H. Viridissima* assemblies revealed that the first (and more ancient) activity peak had a longer duration compared to the shorter first activity peaks observed in *H. oligactis* and *H. vulgaris*. In the latter, the first peak was less intense, also. The second *maT-1* activity peaks in *H. oligactis* or *H. vulgaris* were similar to the first one in their intensity and duration, while that peak was significantly shorter, but more intense in the two representatives of *H. viridissima* (Fig. 6). Meanwhile, some degree of correlation between the TE evolutionary dynamics plots and the TE copy numbers is observed.

Evolutionary dynamics of *maT-2* elements also demonstrates two activity peaks (with the exception of *H. viridissima* assembly (GCA_004118115), in which we could not identify this element with the use of tBLASTn). This element had a single activity peak dated between 11 and 25 MYA, only. In addition, a very low contribution of *maT-2* to the genome of this organism was observed (Fig. 6). As far, as the remaining three assemblies are concerned, some degree of correlation between copy number and genetic coverage, as well, as between the intensity of the recent peaks and the presence of complete and potentially functional TE copies is seen (Fig. 6). tBLASTn search revealed *maT-3* copies in two genomic assemblies of *H. vulgaris* and *H. viridissima* (GCA_004118115), only. Again, one can observe the correlation between total and full-size copy numbers with evolutionary dynamics plots (Fig. 6). A very high recent activity peak found in *H. viridissima* (GCA_004118115) corroborates our hypothesis about potential functionality of *maT-3_HVir/GCA_004118115* element. The presence of *maT-3* elements in *H. oligactis* and *H. viridissima* assemblies (GCA_014706445) was also discovered, however, these TEs demonstrated low genetic coverage and represented only the ancient TE fractions.

Evolutionary dynamics of *maT-4* elements revealed the patterns similar to the previously described three TE types (Fig. 6). *maT-4_HVul*, which we consider to represent a potentially functional copy, demonstrated a high peak of recent activity. *maT-4* could have been detected in *H. oligactis* genome, although with incredibly low genetic coverage.

### Intraspecific differences of *maT* transposons

Due to the availability of two genomic assemblies for two Cnidaria species (*C. quinquecirrha* and *H. viridissima*) in NCBI databases, we had an opportunity to study the intraspecific TE diversities by using the data on TE prevalence, potential functionality and evolutionary dynamics in different representatives of these species (Tab. 1).

**Table 1.**
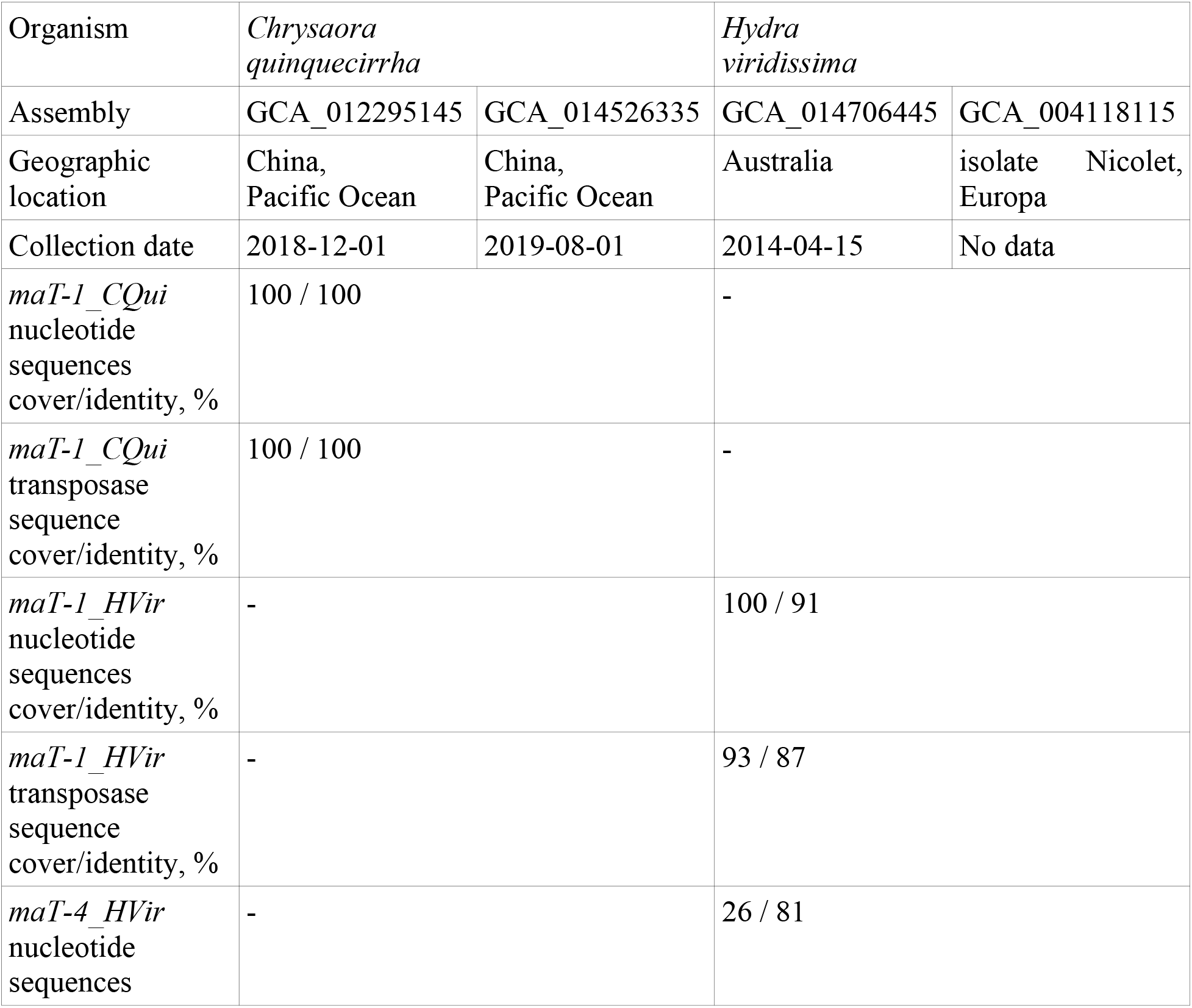
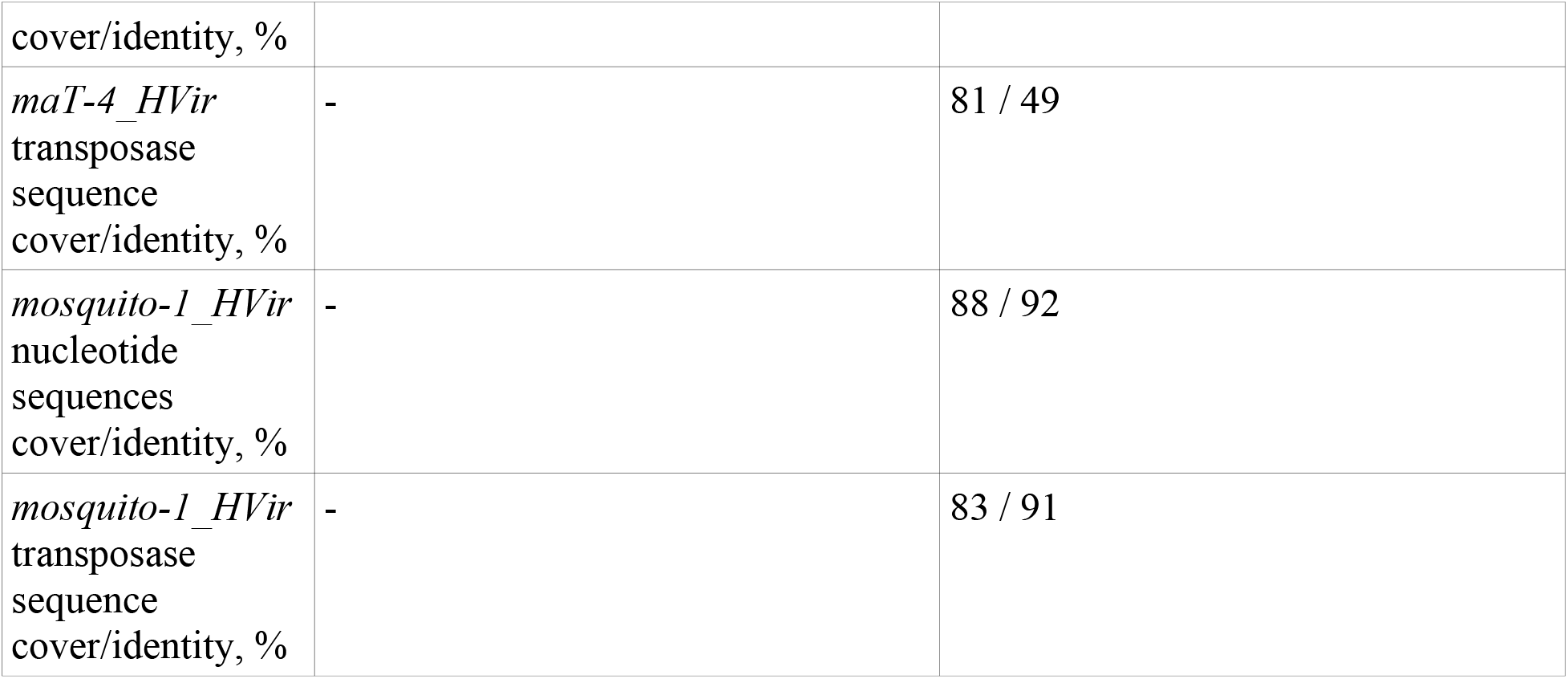
Intraspecific differences of *maT* transposons

The specimens of *C. quinquecirrha* were collected in the Pacifics by a single Chinese research group, which were obtained within 8 months from each other. Most possibly, the individuals belong to the same population. The elements discovered in both assemblies share 100 % identity (Tab. 1). The evolutionary dynamics patterns reconstructed from each assembly were also closely resembling each other (Fig. 5, 6). Such a high sequence identity is not typical for the TEs, which possess high degree of variability.

In case of *H. viridissima*, the first specimen (GCA_014706445) was collected in April, 2014 in Australia, while the second one (GCA_004118115) was a representative of a laboratory stock (strain) kept in Europe. Both assemblies contain *maT-1* and *maT-4* elements (Supplemental Material 1). *maT-1_HVir/GCA_014706445* and *maT-1_HVir/GCA_004118115* elements share high degree of identity of nucleotide sequence (91 %) and the amino acid sequence of transposase ORF (87 %). Both assemblies (GCA_014706445 and GCA_004118115) harbor 11 and 24 complete TE copies, respectively, however, only the latter, but not the former, contain the potentially functional copies. Thus, *maT-1_HVir/GCA_004118115* element, probably, remains active until now, while *maT-1_HVir/GCA_014706445* is being eliminated from the genome of *H. viridissima. maT-4_HVir* element is non-functional in both assemblies and,thus, it is, apparently, completing its life cycle in *H. viridissima*. The patterns of evolutionary dynamics of *maT-1* and *maT-4* are also different (Fig. 6).

In *H. viridissima, maT-3*, which is potentially functional and active element according to GCA_004118115 assembly (with high total and complete copy numbers of 185 and 37 copies, respectively), is completely lacking from GCA_014706445. To the contrary, *maT-2* element is lacking from GCA_004118115, while it is potentially functional and active according to GCA_014706445.

Such an uneven distribution of *maT* elements in different *H. viridissima* assemblies demonstrates a rather high degree of heterogeneity of isolated populations and underscores the diverse ways of TE evolution in different populations. The discovered fact implies that the estimates of TE age and evolution in a particular species require more accuracy, than it can be obtained from the mobilome features of a single individual. Provided the differences in the TE prevalence, copy numbers, and potential functionality, which are that significant between the populations (representing the same species), the interspecific differences can be even more pronounced and, thus, should be taken into account when performing such TE studies.

### Evolutionary history of *maT* elements in Cnidaria

Analyzing the data on copy numbers, evolutionary dynamics and potential functionality, as well as the distribution across taxa and phylogenetic analysis of *maT* transposons, one can conclude that several *maT* invasions have occurred in the cnidarians.

*maT* element discovered by ourselves in the representative of Cubozoa has, most possibly, invaded this class independently of the other *maT* elements found in the cnidarians. Class Cubozoa have diverged about 536 MYA (http://www.timetree.org/). Among the representatives of Cubozoa, we were able to identify *maT* elements in a single family Alatinidae belonging to order Carybdeida, only (Supplemental Material 1). The lack of data does not enable one to suggest when different families have diverged.

The genomic assembly of *Alatinidae sp*. contains only eight seriously damaged *maT-1_ASp* copies and lacks the complete copies. Thus, today *maT* elements are almost completely eliminated from the genome of *Alatinidae sp*. At which stage the invasion of *maT* elements have occurred, can hardly be determined. Whether this invasion have affected only class Cubazoa or it have involved the entire type Cnidaria and have become eliminated from all representatives, except from Cubozoa, also remains unknown. The only fact that can be suggested with a good degree of confidence is that the invasion of this element have occurred very long ago, since *Alatinidae sp* genome harbors the fragments of *maT-1_Asp*, only.

The confirmation that this element was an independent invasion comes from the localization of *maT-1_ASp* on phylogenetic tree, in cluster B1, which does not contain any other *maT* transposon of the cnidarians.

In class Scyphozoa, *maT* elements are found in two families (Ulmaridiae and Pelagiidae) of order Semaeostomeae. Class Scyphozoa have diverged 581 MYA (http://www.timetree.org/), while the divergence of families Ulmaridae and Pelagiidae occured 451 MYA. A comparison of the taxonomic distribution with phylogenetic tree and *maT* element prevalence in Scyphozoa, one can suggest that two invasions of *maT* elements occurred in this class. On phylogenetic tree, *maT* elements found in family Pelagiidae form a separate cluster F with high bootstrap value, while *maT* element found in family Ulmaridae (*Aurelia coerulea*) belongs to cluster A1, which forms a polytomic group together with clusters D, C, and B1, but is only distantly related to cluster F. Among the Ulmaridae, only the genome of *Aurelia coerulea* contains *maT-1_ACoe* element, which has been seriously degraded and is represented by only two strongly damaged copies. The *maT-1_ACoe* invasion have most possibly occurred after the divergence of Ulmaridae и Pelagiidae, since this TE is not present in the latter family. Moreover, the invasion of Ulmaridae by *maT-1_ACoe* is, possibly, an earlier event, compared with the invasion of family Pelagiidae. The genomes of the representatives of Pelagiidae contain multiple *maT* copies (up to 376) while one of elements, *maT-1_CQui/GCA_012295145*, is represented by the fullsize, although not potentially functional copies in the genome of *C. quinquecirrha* (Supplemental Material 1). The degradation of *maT-1_ACoe* element may be either due to its reduced original activity or the significantly earlier time of invasion, which, thus, led to its elimination from *A. coerulea* genome by the present day.

In class Hydrozoa, *maT* elements are found in a single suborder, Aplanulata. The ancestral order, Anthoathecata, have diverged from order Leptothecata 480 MYA (http://www.timetree.org/). Thus, the TE insertions have occurred not earlier, than 480 MYA, since *maT*-elements are found in all representatives of Anthoathecata, while lacking from Leptothecata, but not later, than 193 MYA, since in this period *H. viridissima*, which is the most distantly related species in suborder Aplanulata, have diverged from the common ancestry line. All four *maT* elements found in Hydrozoa are present, in different combinations, in all three hydras (Supplemental Material 1), and all of the four *maT* elements belong to different clusters on phylogenetic tree: *maT-1*, *maT-2*, *maT-3*, and *maT-4* belong to clusters D, C, B2, and A2. We, therefore, suggest that four independent *maT* insertions occurred in Hydrozoa.

Interesting results were obtained when we studied the prevalence and evolutionary dynamics of those four *maT* elements. *maT-1* was found in all species studied. In *H. oligactis*, the full-size and potentially functional copies were not found, although the total copy number was considerably high. To the contrary, the total number of copies present in *H. vulgaris* is not high, but four full-size copies were retained. In *H. viridissima*, eleven full-size elements are found in GCA_014706445 assembly, although none of them is functional, while *maT-1_HVir/GCA_004118115* element from assembly GCA_004118115 can be active, as suggested by the high total and full-size copy numbers and, moreover, two copies are potentially functional ones.

*maT-3* was found in *H. vulgaris* and in GCA_004118115 assembly of *H. viridissima* genome. In the former, the element is represented by a moderate number of copies, five of which are full-size elements, but none of them are potentially functional. *maT-3_HVir/GCA*_004118115 element from *H. viridissima*, to the contrary, shows the features of activity, such as the high total and full-size copy numbers and the three retained potentially functional copies.

We found *maT-4* element in *H. vulgaris* and *H. viridissima*. In the latter, *maT-4* is, presumably, undergoing the elimination stage, while in the former it can be active, since it possesses multiple full-size and four potentially functional copies.

Here, the difference between the estimates of TE insertion time based on evolutionary dynamics analysis or the analysis of TE phylogenies and prevalence should be noted. The time of *maT* insertions in the genomes of hydras estimated by these two approaches is 20-25 MYA or no later, than 193 MYA, respectively. Apparently, the use of neutral mutation rate of 10^−8^/site/year is rather inaccurate for this taxon and, probably, it should be reduced nearly 10-fold.

All *maT* elements described above derive from a common ancestor, since their insertion occurred before the divergence of the hydras. Then, why the same element demonstrates the features of activity (possesses multiple copies) or, otherwise, the degradation in different host species? Apparently, the TE evolution after the divergence of host species occurs differently. The TEs are suggested to undergo neutral evolution (Bourque et al. 2018), and, thus, the patterns of prevalence and evolutionary dynamics can be rather different in different host species or even the populations.

Probably, the host genome and the inserted element interact in an individual manner in every individual. For example, the TE demonstrating an exaggerated activity can be repressed by the genome more aggressively, or the excessive TE transpositions may kill more individuals in the population, while its members with less TE copies gain the advantage and survive. The less active TEs or those that transpose mainly into the non-coding regions of the genome and do not affect the essential genes can preserve their activity and remain in the host genomes for a longer period of time (Casacuberta and González 2013). In isolated cases, the possibility of the TE life cycle restart can not be ruled out. The TE reactivations were shown to possibly occur due to inbreeding or outbreeding Speciation is directly linked to the processes of inbreeding (bottleneck) and outbreeding (population merge). The secondary (repeated) outburst of TE activity (life cycle restart) leading to the increase in their copy numbers in particular species can occur via these mechanisms.

### *mosquito* elements in Cnidaria

In our work, we identified the elements of representatives of *mosquito* (DD37E) family in the representatives of Hydrozoa (Aplanulata family), namely, in *H. viridissima*, *H. oligactis* and *H. vulgaris*. This TE family remains understudied. Previously, the discovery of the *mosquito* family TEs in insects and the ctenophores was published (Shao and Tu 2001; Wang et al. 2021), however, the prevalence of *mosquito* TEs among the cnidarians was not studied and reported. Using the sequence of the previously discovered *mosquito* elements as a query, we further investigated those genomic assemblies of the cnidarians available from NCBI, however, the *mosquito* family representatives have not been identified elsewhere, except in class Hydrozoa.

Only a single element, which we refer to as *mosquito-1*, is found in all species of the hydrozoans. The potentially functional copies of the TE were present in GCA_014706445 genomic assembly of *H. viridissima*. This element has the length of 1309 bp, an intact transposase ORF of 341 amino acid residues, and 31 bp TIRs. The total and full-size copy numbers are 162 and 65, respectively, which indicates at a recent high activity of this TE in the genome of *H. viridissima* (Supplemental Material 1). The three genomic assemblies contained the damaged *mosquito* copies only. Their length varied between 702 and 1445 bp, the size of transposase ORF ranged from 233 to 406 aa residues. The described element, *mosquito-1_HOli*, possessed TIRs of 30 bp in length (Supplemental Material 1). The total copy numbers of the nonfunctional elements in the genomes of hydras, was rather high (149, 279 and 349 copies), which indicates at the past activity of these TEs, which has not been retained until currently, as judged by the presence of their damaged copies, only.

We analyzed the structure of *mosquito-1_HVir/GCA_014706445* transposase protein in order to probe its potential functionality. It possesses the catalytic domain typical for *mosquito* family, which contain DD37E, PAIRED domain with 6 well demarcated alpha helices, and GRPR-like motif and NLS sequence with their typical localization (Fig. 2). Based on the described features, two *mosquito-1_HVir/GCA_014706445* copies can be considered as the potentially functional ones.

An analysis of the evolutionary dynamics of *mosquito-1* elements revealed their differences in patterns and insertion time. The maximum value of statistical parameter k never exceeded 25% for *mosquito-1_HOli*, while in the rest three assemblies it varied from 44 to 50% (Fig. 7). Moreover, *mosquito-1_HOli* had only a single peak of activity, while the other three had 2 peaks. The results of phylogenetic analysis reveal that all *mosquito* elements discovered by ourselves form a single clade with the previously described *mosquito* elements (Shao and Tu 2001; Wang et al. 2021). Taking into acount the data provided by TimeTree (http://www.timetree.org/), we suppose that the invasion of *mosquito* elements occurred between 480 and 193 MYA, before the speciation event, since they are present in Hydrozoa only. A comparison of transposon distribution between two *H. viridissima* assemblies revealed rather contrast differencies (Supplemental Material 1). As mentioned before, the individual *H. viridissima* specimens, from which GCA_014706445 and GCA_004118115 assemblies were derived, have been collected from different populations. A comparison of their elements yielded a high level of sequence identity, 91,9 % and 90,8 % for nucleotide and amino acid sequence, respectively (Supplemental Material 1). However, a comparison of the element representation in these two individuals shows that *mosquito-1_HVir/GCA_014706445* is active, since it is represented in multiple copies including the full-size ones and those that can potentially remain functional. Meanwhile, *mosquito-1_HVir/GCA_004118115* element is represented in the genome by the damaged copies, although these are multiple, which indicates at its activity in the past. The patterns of evolutionary dynamics are also different for the two elements. An ancient activity peak is more intense and long in GCA_004118115 assembly, while in GCA_014706445 assembly the reverse pattern with a broader recent peak is observed, which correlates with the presence of the full-size copies.

**Fig. 7.**
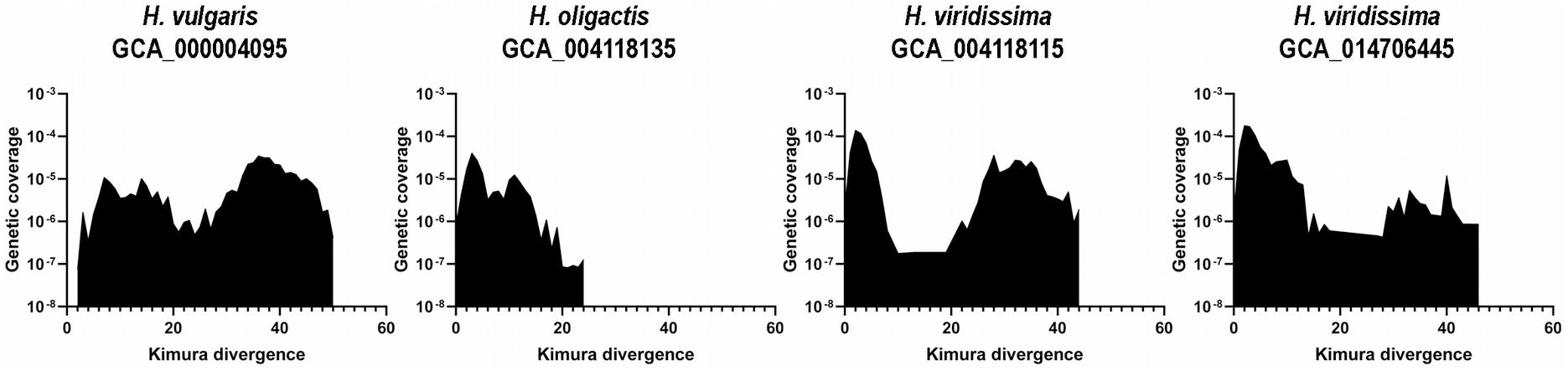
Evolutionary dynamics of hydras *mosquito* transposons. The X-axis indicates the Kimura divergence estimate (%), the Y-axis represents the log10 of copies coverage (%) in the genome.

These intraspecific differences again corroborate the conclusion made for *maT* elements. Despite the TE evolution is thought to be neutral, the TEs exert strong influence on the genomes and can evolve in an unpredictable manner. The populations that have diverge recently in the course of evolution show contrasting difference of the evolution history of two deifferent elements, *maT* and *mosquito*. Thus, any element inserted in the genome and remaining active there induces a completely unique host genome response leading either to elimination or to proliferation of the element. The mechanisms of element containment have been previously described, although they still remain understudied. The lack of correlation between the laboratory and the wild populations is an intriguing fact.

## Author Contributions

Puzakov M. conceived and designed the study. Puzakov M., Puzakova L. and Shi S. mined transposons, collected data and performed the data analysis. Puzakov M. and Cheresiz S. wrote the manuscript. All authors have read and approved the final version of the manuscript.

## Supplemental Materials

Supplemental Material 1. Cnidarian DD37E/D transposons (TIR – terminal inverted repeats; SIR – subterminal inverted repeats; bp – base pairs; aa – amino acids; NI - not identified)

